# Immune Classification of Clear Cell Renal Cell Carcinoma

**DOI:** 10.1101/2020.07.03.187047

**Authors:** Sumeyye Su, Shaya Akbarinejad, Leili Shahriyari

## Abstract

Since the outcome of treatments, particularly immunotherapeutic interventions, depends on the tumor immune micro-environment (TIM), several experimental and computational tools such as flow cytometry, immunohistochemistry, and digital cytometry have been developed and utilized to classify TIM variations. In this project, we identify immune pattern of clear cell renal cell carcinomas (ccRCC) by estimating the percentage of each immune cell type in 526 renal tumors using the new powerful technique of digital cytometry. The results, which are in agreement with the results of a large-scale mass cytometry analysis, show that the most frequent immune cell types in ccRCC tumors are CD8+ T-cells, macrophages, and CD4+ T-cells. Saliently, unsupervised clustering of ccRCC primary tumors based on their relative number of immune cells indicates the existence of four distinct groups of ccRCC tumors. Tumors in the first group consist of approximately the same numbers of CD8+ T-cells, CD4+ T-cells, and macrophages, while tumors in the second group have a significantly high number of macrophages compared to any other immune cell type (P-value< 0.01). The third group of ccRCC tumors have a significantly higher number of CD8+ T-cells than any other immune cell type (P-value< 0.01), while tumors in the group 4 have approximately the same numbers of macrophages and CD4+ T-cells and a significantly smaller number of CD8+ T-cells than CD4+ T-cells (P-value< 0.01). Moreover, there is a high positive correlation between the expression levels of IFNG and PDCD1 and the percentage of CD8+ T-cells, and higher stage and grade of tumors have a substantially higher percentage of CD8+ T-cells. Furthermore, the primary tumors of patients, who are tumor free at the last time of follow up, have a significantly higher percentage of mast cells (P-value< 0.01) compared to the patients with tumors for all groups of tumors except group 3.

## Introduction

Clear cell renal cell carcinoma (ccRCC) is the most frequently diagnosed malignant tumor type in the adult kidneys consisting of approximately 85% of kidney cancer cases^1^, and surgical resection is the common therapy type for ccRCC. However, it is not effective for patients with advance or metastatic ccRCC^2^. Several immunotherapeutic approaches have been recently used for treating patients with ccRCC^3, 4^, which is considered a morphologically and genetically immunogenic tumor^5^. However, many patients do not respond to these treatments and develop adaptive or intrinsic resistance. We can increase the response rate to these treatments by identifying types of tumors that would respond to them.

Several studies show that cancer cells and tumor infiltrating immune cells (TIICs), which have important roles in both regulation of cancer progression and promotion of tumor development^6, 7^, play an important role in the determination of malignant tumor types^8, 9^. Tumor-infiltrating lymphocytes (TILs), which include T-cells and B cells, are an important category of TICCs. CD4+ helper T-cells and cytotoxic CD8+ T-cells play a significant role in preventing tumor by targeting antigenic tumor cells^10^, and CD8+ T-cells are linked with better clinical outcomes and reaction to immunotherapy in many cancers^11, 12^. Furthermore, it has been recently observed that tumor associated B cells, which have significant roles in the immune system by producing antibodies and presenting antigens, could be predictors of survival and response to immune checkpoint blockade therapy^13^. Additionally, a relationship between TIICs gene signatures and lower survival rates has been observed in ccRCC patients, and tumor-associated macrophages (TAM) and 22 T cell phenotypes are found to be correlated with clinical outcomes^14, 15^. These observations emphasize on importance of analyzing the cellular heterogeneity of tumors, including immune cell variations, to identify target tumors for each specific treatment and design new effective cancer treatments^16^.

There are some experimental approaches such as single cell analysis tools, including immunohistochemistry and flow cytometry to observe tumor immune infiltrates, however these methods are expensive and time consuming, and they are limited to analyzing a few immune cell types simultaneously^17^. For this reason, several computational methods have been recently developed to provide us with much less expensive and fast alternative ways to estimate the relative amount of each cell type from gene expression profiles of bulk tumors. In this study, we applied a powerful “digital cytometry” method called CIBERSORTx^18^ to determine immune patterns of tumors (Figure 1A) and investigate the association of these patterns with clinical features.

**Figure 1.**
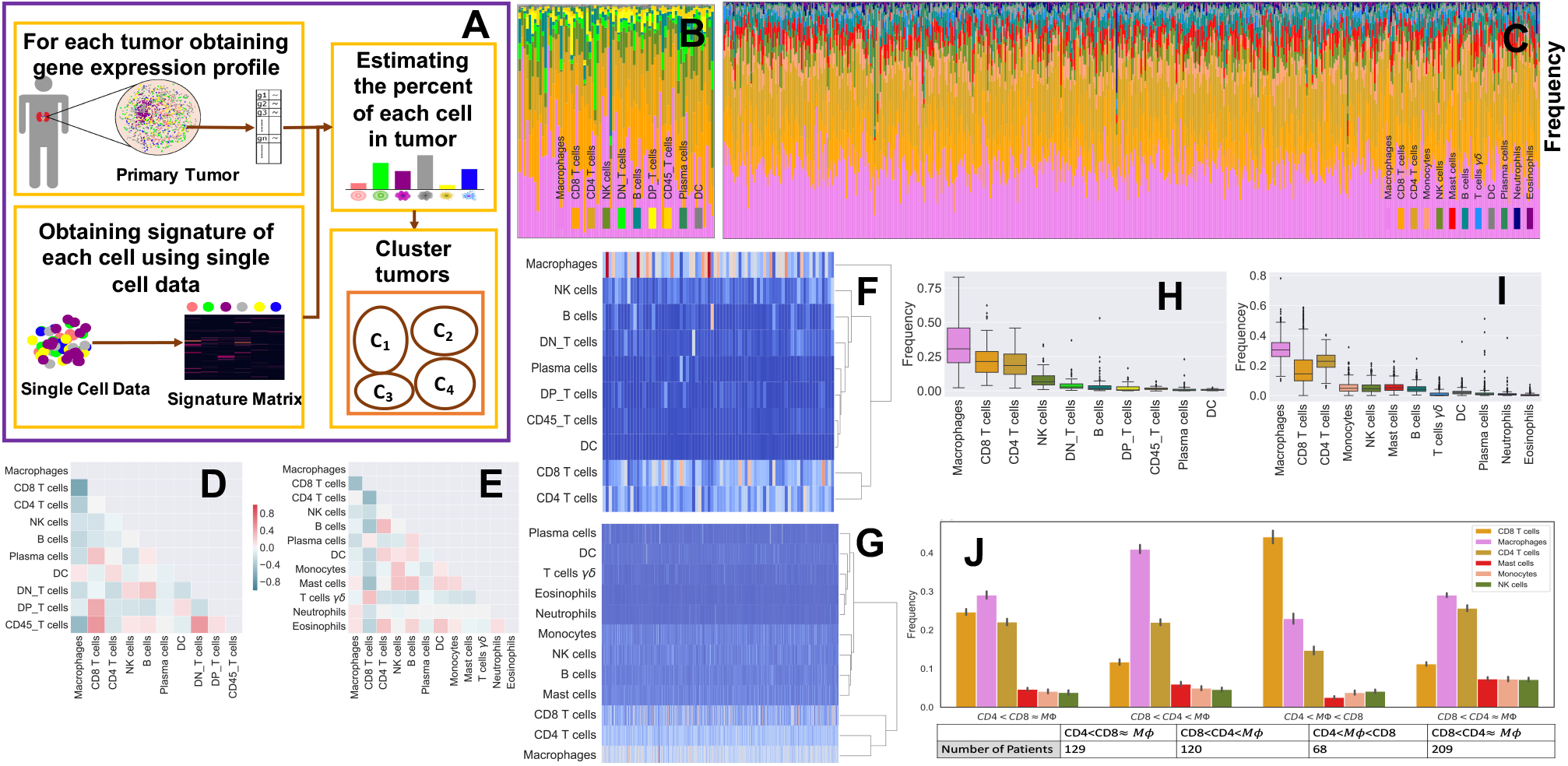
Immune pattern of ccRCC. Sub-figure A represents the algorithm of digital cytometry and clustering applied on TCGA data. Sub-figures B and C respectively show the estimated percentage of each immune cell by mass cytometry analysis of 73 ccRCC patients done by Chevrier et al^14^ (B) and digital cytometry on 526 TCGA ccRCC tumors (C). Sub-figures D and E indicate the correlation map of estimated immune cell frequencies in 73 ccRCC tumors (D) and TCGA ccRCC tumors (E), respectively. Sub-figures F and G show the cluster heat map of immune cell frequencies in 73 ccRCC tumors (F) and TCGA ccRCC tumors (G). Sub-figures H and I respectively show a box plot format of the immune cell percentages in 73 ccRCC tumors (H) and TCGA ccRCC tumors (I). Sub-figure J shows 4 distinct immune patterns of ccRCC tumors obtained by K-mean clustering of cell frequencies of TCGA ccRCC tumors.

## Results

To estimate the percentage of each cell type in ccRCC tumors, we apply “digital cytometry”, CIBERSORTx, on TCGA gene expression profiles of ccRCC primary tumors. We compare the results of our “digital cytometry” analysis with the results of an experimental study of a large-scale mass cytometry-based immune cells analysis of 73 ccRCC patients^14^. Immune cells, which have been characterized in this experimental study done by Chevrier et al.^14^ are macrophages, CD8+ T-cells, CD4+ T-cells, NK cells, B cells, plasma cells, dendritic cells (DC), CD45+ T-cells, double positive T-cells (DP_T-cells), double negative T-cells (DN_T-cells). To be able to compare our results, which includes 22 immune cell types given in LM22 signature matrix of CIBERSORTx, we combine cells that belong to the same family. For instance, since CD4+ naive T-cell, CD4+ memory resting T-cells, CD4+ memory activated T-cells, follicular helper T-cells, and regulatory T-cells are sub-types of CD4+ T-cells, we sum their numbers to estimate the total number of CD4+ T-cells. We do similar calculation for B cells, NK cells, DC cells, macrophages, and mast cells.

### The most frequent immune cells in ccRCC tumors are macrophages, CD4+ T-cells, and CD8+ T-cells

Results of experimental study done by Chevrier et al^14^ show that macrophages are the most frequent immune cells in most ccRCC tumors with a mean of 31% followed by CD8+ T-cells and CD4+ T-cells, respectively (H), which are in agreement with the results of CIBERSORTx applied on TCGA data set (Figure 1C,I).

### There is a negative correlation between the number of macrophages and CD8+ T-cellsd

The results of mass cytometry analysis indicate a negative correlation between CD8+ T-cells and macrophages with Pearson correlation coefficients of −0.67. Importantly, the digital cytometry applied on TCGA data set confirms this negative correlation between the number of CD8+ T-cells and macrophages in ccRCC with a correlation coefficient of −0.46 (Figure 1D,E).

### Variations of ccRCC tumors are mainly in the percentage of macrophages, CD8+ T-cells, and CD4+ T-cells compared to the other immune cell types

Figure 1 shows high variations among the percentage of CD8+ and CD4+ T-cells and macrophages across ccRCC tumors, while there is a slight variation in the percentage of other immune cell types. Unsupervised hierarchical clustering of cell frequencies show that CD8+ T-cells and CD4+ T-cells are clustered together in the experimental results, and then they group with macrophages and other cells (Figure 1F). The result of digital cytometry on TCGA data shows a kind of similar trend: CD4+ T-cells first clustered with macrophages, then they clustered with CD8+ T-cells and other cells (Figure 1G).

### There are four immune patterns of ccRCCs

K-mean clustering of ccRCC tumors based on their immune cells’ frequencies shows that there are four different immune classes: Cluster 1 (*CD*4 < *CD*8 ≈ *M*Φ), in which the numbers of macrophages and CD8+ T-cells are approximately the same, and the number of CD4+ T-cells is slightly less than the number of CD8+ T-cells; Cluster 2 called (*CD*8 < *CD*4 < *M*Φ), in which the number of macrophages is significantly higher than the number of CD4+ and CD8+ T-cells (P-value< 0.01); Cluster 3 (*CD*4 < *M*Φ < *CD*8), in which the number of CD8+ T-cells is significantly higher than the number of macrophages and CD4+ T-cells (P-value< 0.01); and Cluster 4 called (*CD*8 < *CD*4 ≈ *M*Φ) in which the numbers of macrophages and CD4+ T-cells are approximately the same, and the number of CD8+ T-cells is significantly less than CD4+ T-cells (P-value< 0.01) (Figure 1J).

### Cluster (*CD*8 < *CD*4 ≈ *M*Φ) has the highest percentage of grade and stage 1 and 2 tumors

Comparing clinical features of clusters show that Cluster (*CD*8 < *CD*4 ≈ *M*Φ) includes the highest percentage of grade 1 and grade 2 tumors and the lowest percentage of grade 4 tumors, and there is a similar trend for the stage of tumors (Figure 2A,B). Importantly, clusters (*CD*8 < *CD*4 ≈ *M*Φ) and (*CD*8 < *CD*4 < *M*Φ) have the highest proportion of patients who were tumor free and smallest percentage of the diseased patients at the last time of follow up among all other clusters (Figure 2C,D). Furthermore, this cluster has the highest frequency of mast cells, monocytes and B cells compared the other clusters (Figure 1J). These results might imply that non-aggressive tumors include an approximately equal number of each immune cell type.

**Figure 2.**
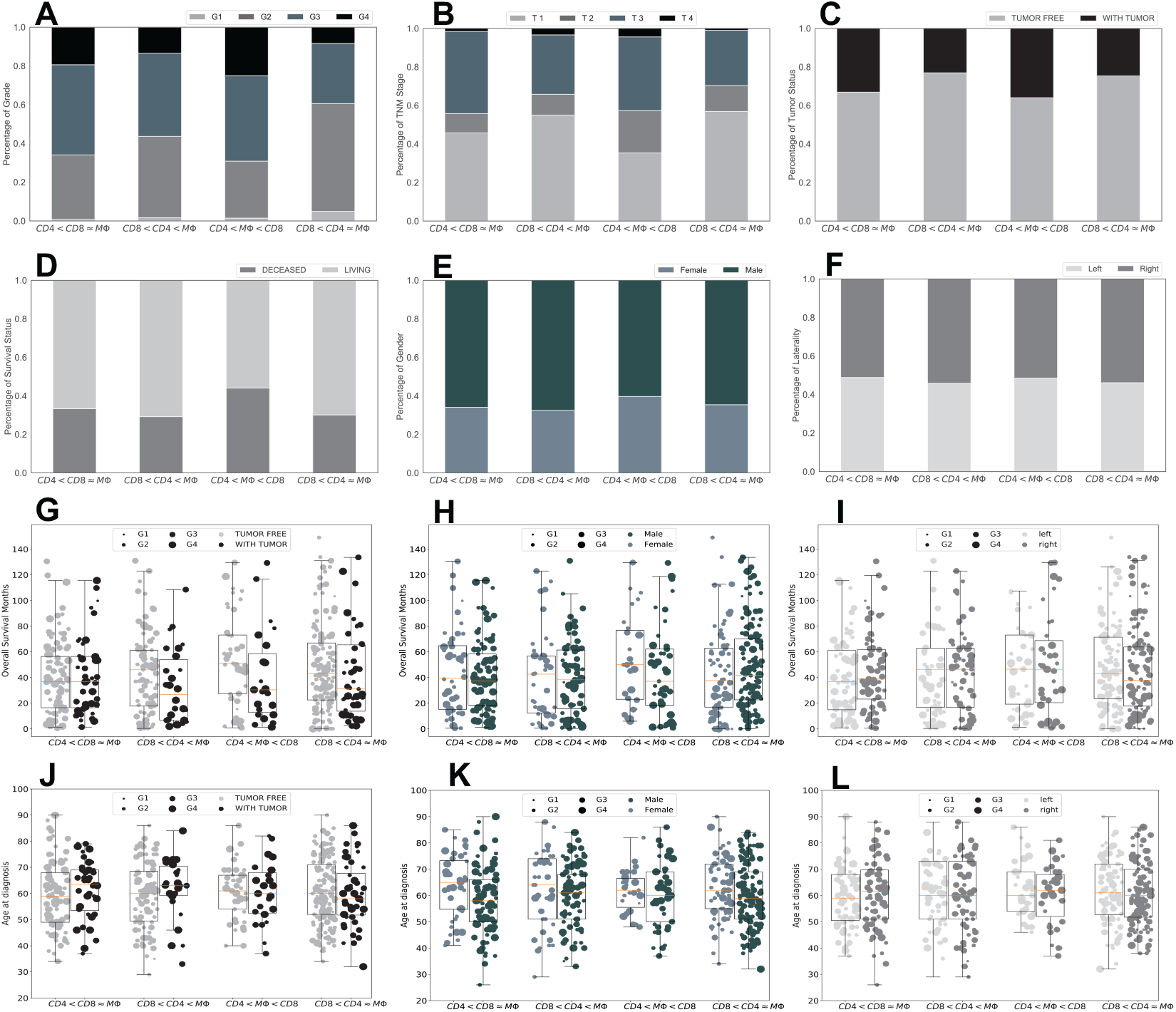
Clinical features of each ccRCC tumor cluster. Sub-figures A-F show the percentage of patients with grade 1-4 (A), stage 1-4 (B), with tumors or without tumors (C), alive or dead at the last time of follow up (D), female or male (E), and primary tumors in left or right kidney (F) for each cluster of ccRCC tumors. Sub-figures G-I and J-L respectively show the overall survival months and age of diagnosis of the patients in each cluster as a function of tumor status (G,J), gender (H,K), and the location of the primary tumor (I,L); the size of markers indicates the grade of tumors.

### Cluster (*CD*4 < *M*Φ < *CD*8) has the highest percentage of grade and stage 4 tumors compared to the other clusters

The percentages of grade and stage 3 and 4 tumors are higher in Cluster (*CD*4 < *M*Φ < *CD*8) compared to the other clusters (Figure 2A,B). Furthermore, this cluster includes the highest number of deceased patients and patients who had a tumor at the last time of follow up compared to the other clusters (Figure 2C,D). There is a noticeable difference among overall survival months of female and male patients in this cluster, female patients in the cluster (*CD*4 < *M*Φ < *CD*8) have the highest overall survival months compared to the other clusters (Figure 2H). These results indicate that male patients’ ccRCC tumors consisting of a significantly higher number of CD8+ T-cells than any other immune cell types might be aggressive.

### There is no significant differences in overall survival months or age at diagnosis between clusters

Figure 2 indicates no significant differences in the overall survival of patients between any of these clusters; this figure also reveals some other interesting observations. For example, patients in Cluster *CD*4 < *CD*8 ≈ *M*Φ with and without tumors at the last time of follow up have a similar overall survival months while in all other clusters patients with tumor have a substantially lower survival months than patients without tumors at the last time of follow up (Figure 2G). Moreover, patients with tumor in this cluster have a remarkably higher age at diagnosis compared to the patients with no tumors in this cluster (Figure 2J). Furthermore, female patients in this cluster have a noticeably higher age at diagnosis but the same survival as male patients in this cluster (Figure 2H,K). Additionally, female patients in Cluster *CD*4 < *M*Φ < *CD*8 have a substantially higher overall survival months than male patients in this cluster, while females have a slightly higher age at diagnosis than males in this cluster. Importantly, there is no significant differences in the age at diagnosis and survival months of patients in each cluster based on the location of their primary tumors, left and right kidneys (Figure 2I,L).

### Higher grade and stage of ccRCC tumors have higher percentage of CD8+ T-cells and lower percentages of mast cells and monocytes

A study of 87 ccRCC patients indicates that the percentage of tumor infiltrating CD8+ T-cells co-expressing PD-1 and Tim-3 correlated with an aggressive phenotype and a larger tumor size at diagnosis^19^. Figure 3 also shows that the grade and stage 3 and 4 ccRCC tumors have a significantly higher percentage of CD8+ T-cells compared to the stage and grade 1 and 2 tumors (P-value< 0.01), which is consistent with the observations of Figure 2.

**Figure 3.**
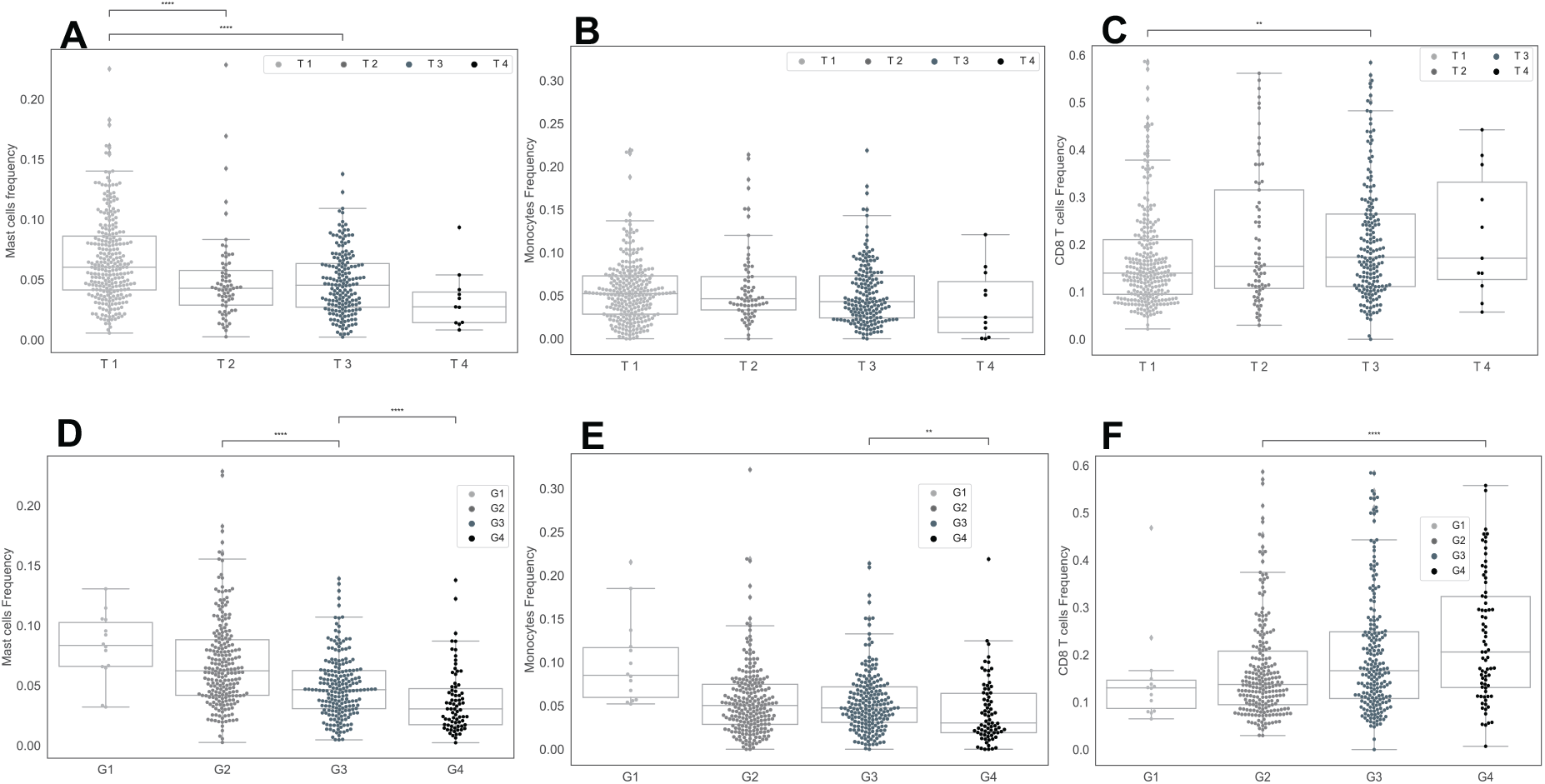
Percentage of mast cells, monocytes and CD8+ T-cells in ccRCC tumors as a function of grade and TNM staging. Sub-figures A-C show the percentages of mast cells (A), monocytes (B), and CD8+ T-cells (C) in primary tumors as a function of stage of tumors. Sub-figures D-F represent the percentage of mast cells (D), monocytes (E), and CD8+ T-cells (F) in primary tumors as functions of grade of tumors.

Figure 3 also indicates that the percentages of mast cells and monocytes in ccRCC tumors significantly decrease when the grade and stage of tumors increase (P-value< 0.01). Note, Clusters (*CD*8 < *CD*4 < ≈ *M*Φ) and (*CD*8 < *CD*4 ≈ *M*Φ) that have higher frequency of mast cells and monocytes and lower frequency of CD8+ T-cells have the least percentage of grade three and four tumors (Figs 1J and 2).

### Tumor free patients have a significantly higher percentages of mast cells in their primary tumors

Figure 4A shows that primary tumors of patients who are tumor free at the last time of follow up has a significantly higher level of NK cells compared to the patients with tumor ((P-value< 0.01). However, a closer look in clusters reveal that the significant difference (P-value< 0.01) in percentage of NK cells between tumor free and with tumor patients corresponds to the patients in Cluster (*CD*4 < *CD*8 ≈ *M*Φ) (Figure 4B). However, in all clusters, the percentage of mast cells is higher in primary tumors of tumor free patients versus with tumor patients at the last time of follow up. Importantly, Cluster (*CD*8 < *CD*4 ≈ *M*Φ) has the highest percentage of mast cells and NK cells compared to the other clusters (Figs 1J and 4). Note, this cluster has the highest percentage of grade and stage 1 and 2 tumors. Additionally, ccRCC tumors in Cluster (*CD*4 < *M*Φ < *CD*8), which has the highest percentage of grade and stage 4 tumors, have the lowest amount of mast cells.

**Figure 4.**
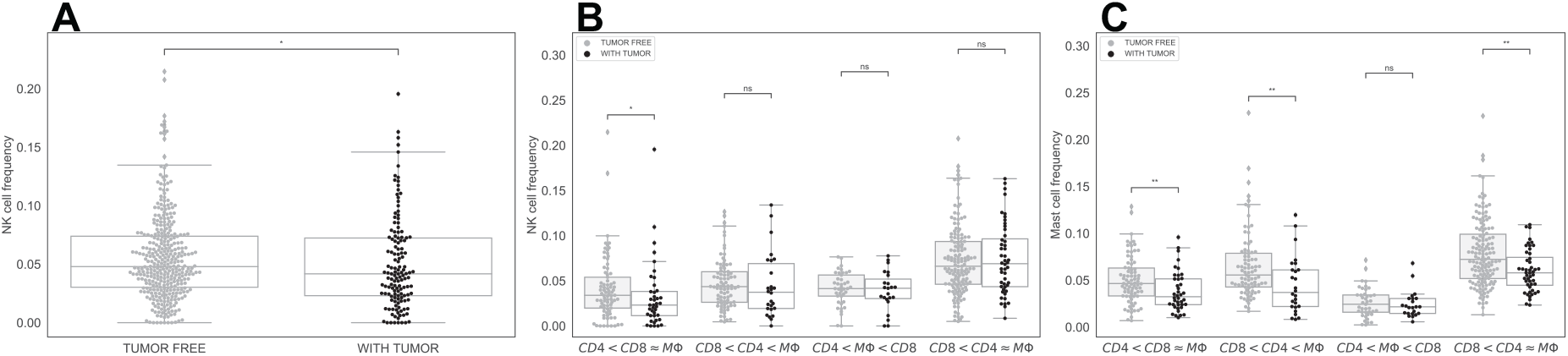
Frequency of NK cells and mast cells in ccRCC. Sub-figure A shows that patients who were tumor free at the last time of follow up have higher percentage of NK cells than patients with tumor at the last time. Sub-figures B and C respectively indicate the percentage of NK cells and mast cells in primary tumors in each cluster.

### Genes expression levels of PDCD1 and INFG are significantly positively correlated with the percentage of CD8+ T-cells in ccRCC tumors

Programmed cell death protein 1 (PD-1) is a type of protein that found on T-cells and it prevents T-cells from killing cancer cells when it binds to PD-1 ligand (PD-L1) and PD-2 ligand (PD-L2) on cancer cells^20^. PDCD1 gene, which encodes PD-1 protein^21^, and CD8+ T-cells are highly positively correlated, with correlation coefficient of 0.85. Also, expression level of PDCD1 is the highest in the cluster (*CD*4 < *M*Φ < *CD*8) and the lowest in the cluster (*CD*8 < *CD*4 ≈ *M*Φ) as a result of positive correlation with CD8+ T-cells (Figure 5C,E).

**Figure 5.**
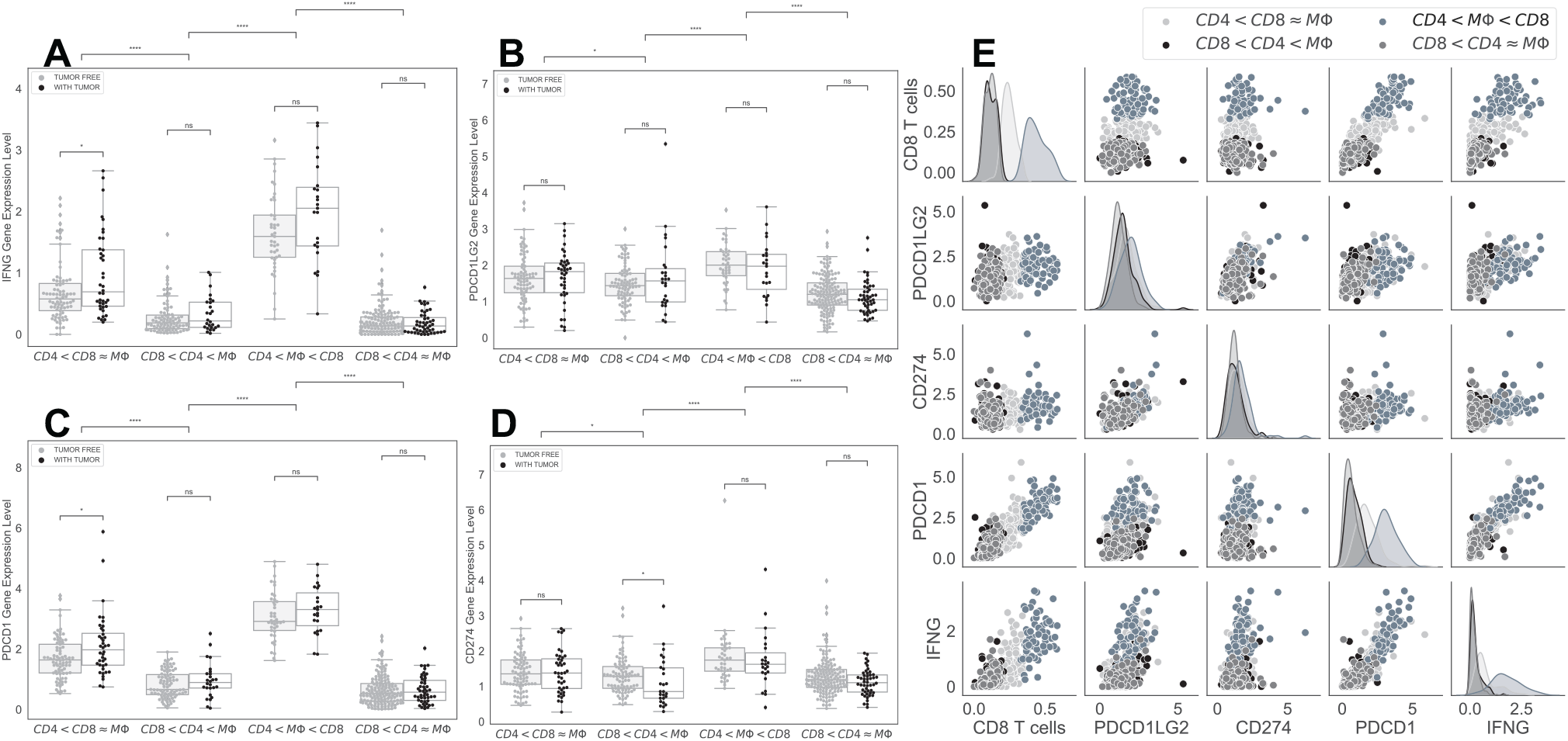
Expression levels of genes encoding PD-1, PD-L1, PD-L2, and IFN *γ*. Sub-figures A-D indicate the expression levels of INFG, PDCD1LG2, PDCD1 and CD274 in each cluster as a function of tumor status, respectively. Sub-figure E represents the correlations and distributions of INFG, PDCD1LG2, PDCD1, CD274 expression levels and CD8+ T-cells; color coded based on the clusters.

Interferon *γ* (*INFγ*), encoded by INFG gene^22^, is a cytokine that is essential for innate and adaptive immunity. It works as an activator of macrophages and stimulator of NK cells and neutrophils^23^, and it is mostly produced by T-cells and NK cells as a reaction of a variety of inflammatory or immune stimuli^24^. Saliently, expression level of INFG is significantly positively correlated with the percentage of CD8+ T-cells and the expression level of PDCD1 in ccRCC tumors, with correlation coefficients of 0.79 and 0.87, respectively. In addition, cluster (*CD*4 < *M*Φ < *CD*8) has the highest INFG expression level and cluster (*CD*8 < *CD*4 ≈ *M*Φ) has the lowest expression level of INFG as expected (Figure 5).

In contrast, there is a slightly positive correlation between the expression levels of CD274 and PDCD1LG2 genes, that encodes PD-L1 and PD-L2 respectively, with the expression levels of PDCD1 and INFG, and the percentage of CD8+ T-cells in ccRCC tumors (Figure 5E). In addition, cluster (*CD*8 < *CD*4 ≈ *M*Φ) has the lowest levels of CD274 and PDCD1LG2 compared to the other clusters (Figure 5B and D).

### There is a significant association between RGS5 expression level and the percentages of NK cells, monocytes, and mast cells

RGS5 is a member of the regulators of G protein signaling (RGS) family, and they are known as signal transaction molecules that are associated with the arrangement of heterotrimetric G proteins by acting as GTPase activators. Moreover, RGS5 is a hypoxia-inducible factor-1 dependent involved in the induction of endothelial apoptosis^25^. In our previous study on TCGA data, we found that a high expression level of RGS5 in ccRCC primary tumors is associated with better survival months, and when the grade of ccRCC tumor increases, the RGS5 expression level significantly decreases^26^. Interestingly, cluster (*CD*8 < *CD*4 ≈ *M*Φ) has the highest RGS5 expression level compared to the other clusters, and tumor free patients have a higher level of RGS5 expression than patients with tumor (Figure 6A). Saliently, ccRCC tumors with a high expression level of RGS5 have a significantly high percentages of NK cells, mast cells, and monocytes (P-value< 0.01) (Figure 6B-D).

**Figure 6.**
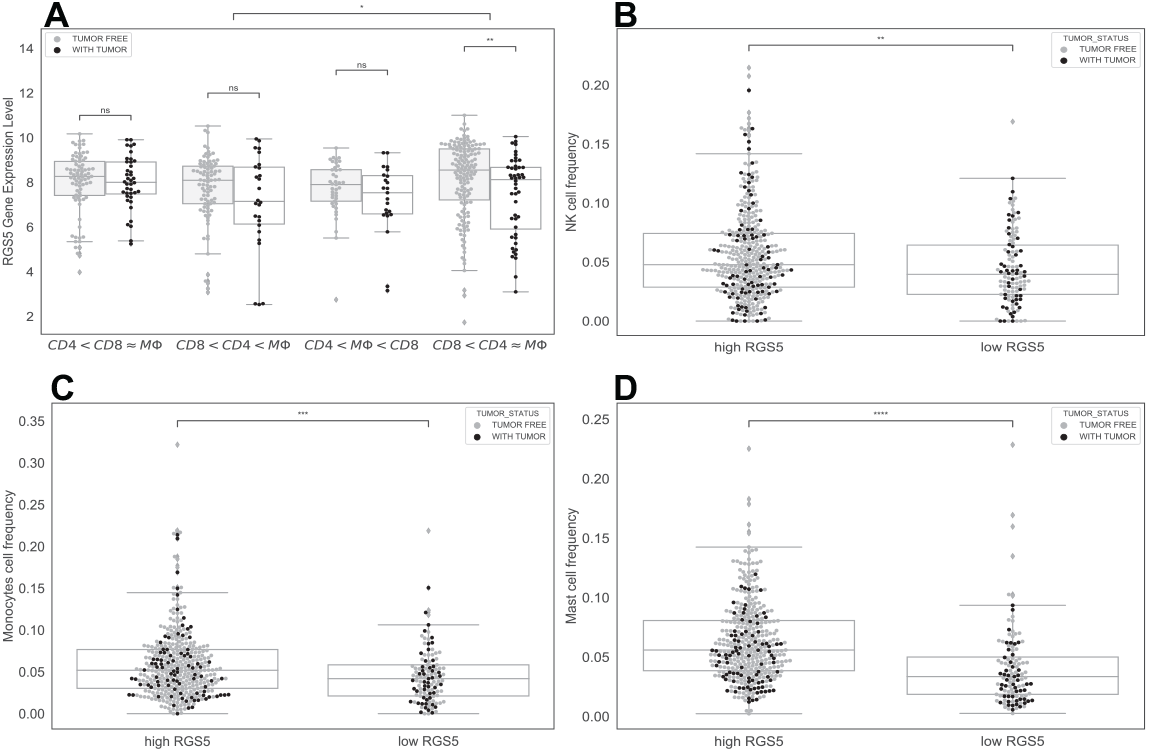
RGS5 expression level in ccRCC tumors. Sub-figure A shows the expression level of RGS5 in ccRCC tumors in each cluster as a function of tumor status. Sub-figures B, C and D indicate the relation between the level of RGS5 and the percentages of NK cells, monocytes, and mast cells in ccRCC tumors, respectively.

## Discussion

Immune checkpoints are essential parts of immune system, and they are crucial to prevent autoimmune diseases. However, some tumors benefit from these checkpoints, because these checkpoints can prevent the immune system from killing cancer cells. One such immune checkpoint is programmed cell death 1 (PD-1) protein, which binds to its ligand PD-L1 and inhibits immune cell activities, including T cell activities. One strategy for cancer immunotherapy is to block these checkpoints to promote anti-cancer T-cell activities^27–30^. Immunotherapy such as targeting PD-1 pathway has improved overall survival months of several patients with metastatic cancers, including melanoma, head and neck cancer, renal cell carcinoma, non-small cell lung cancer (NSCLC), and colon cancer. However, there are many patients who do not respond to these treatments, and some patients who initially respond to the treatments, they might develop resistance or experience severe adverse events^31–33^. For this reason, further biomarkers of tumor cells such as PD-1 and PD-L1 and of tumor infiltrating immune cells such as T-cells and macrophages need to be established to develop new treatment strategies and identify the patients who can be treated by each drug or treatment strategy^34^.

In kidney cancer, common immunotherapy drugs such as nivolumab and avelumab target PD-1, PD-L1, and PD-L2 pathways^35^. Anti PD-1 drugs targets T-cells directly, while anti-PD-L1 drugs target tumor cells directly, and they may also target tumor associated macrophages that express PD-L1. Several studies indicate an increase of *INFγ* production in the PD-1 inhibitors and other immune checkpoint blockade therapies that resulted in destruction of cancer cells^36–38^, and a relation between cancer immunotherapy improvement and an increase of *INFγ* expression has been observed^24^. Moreover, a correlation observed between an increase in *INFγ* gene expression and better progression-free survival in NSCLC and urothelial cancer patients treated with a PD-L1 inhibitor^39^.

Note, tumors in cluster (*CD*4 < *M*Φ < *CD*8) have a high expression levels of INFG, the gene encoding *INFγ*, and PDCD1, the gene encoding PD-1, compared to the other clusters, and the expression levels of these genes are significantly positively correlated with the percentage of CD8+ T-cells in tumors. Importantly, it has been shown that *INFγ* boosts the CD8+ T-cells expansion^40^. Thus, patients in the cluster (*CD*4 < *M*Φ < *CD*8) might respond to the PD-1 inhibitors. In addition, since there is not a strong correlation between PDCD1LG2 and CD274 expression levels and levels of INFG and PDCD1 genes, PD-L1 and PD-L2 inhibitors might not be as effective treatments as the PD-1 inhibitors for the patients in this cluster. Although Cluster (*CD*8 < *CD*4 ≈ *M*Φ) includes a high number of patients with lower grade and without tumor in the last follow up time, tumors in this cluster have lower levels of INFG and PDCD1, therefore patients in this cluster may not be a good candidate for anti PD-1 therapies.

Anti-angiogenic agent (AA) is one of the main treatments in the aggressive ccRCC^1^, because nutrients and oxygen are the main ingredients of the tumor growth which come from blood. Anti-angiogenics, also known angiogenesis inhibitors, are drugs that stop the growth of blood vessels (angiogenesis) that tumors need to grow^41^. A study of in vitro cell lines and in vivo mouse models of ccRCC shows that the recruitment of mast cells is related with increased ccRCC angiogenesis by modulating *PI*3*K* ⟶ *AKT* ⟶ *GSK*3*β* ⟶ *AM* signaling pathway^42^. Since Cluster (*CD*8 < *CD*4 ≈ *M*Φ) has the highest amount of mast cells compared to the other clusters, angiogenesis inhibitors might be a good treatment option for the patients in this cluster. Moreover, mast cells are suggested as an independent prognostic factor in some studies of ccRCC patients^43, 44^, and the amount of mast cells negatively correlated with 5-year survival^44^. However, our finding shows that the number of mast cells inversely correlated with the grade of tumors (Figure 3A, D), and the primary tumors of patients without tumors at the last time of follow up have higher percentages of mast cells than primary tumors of patients with tumor at the last time of follow up.

Kruger et all^45^ suggested RGS5 gene as a tumor associated antigenes (TAAs), and they observed over-expressed RGS5 level from a large scale analysis of ccRCC specimens. Another study found that RGS5 is strongly up-regulated in a broad variety of malignant cells and showed that RGS5 peptides might be a good candidate for designing cancer vaccines to target malignant cells and tumor vessels^46^. We found that patients with higher RGS5 levels have significantly higher percentages of NK cells, mast cells, and monocytes in their primary tumors (P-value< 0.01). Moreover, patients in Cluster (*CD*8 < *CD*4 ≈ *M*Φ) have the highest amount of RGS5 expression in their primary tumor. With the help of further investigation, RGS5 gene might be a good target for patients in this cluster. Further clinical and biological studies are required to test and validate all above mentioned suggestions.

## Materials and methods

We estimated the percentage of tumor infiltrating immune cells in ccRCC tumors using CIBERSORTx deconvolution method that is based on the following linear model:

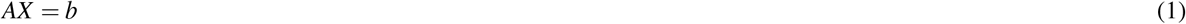

where *b*, which is called mixture data, is the gene expression profile of the bulk tumor, and *X* is unknown cell proportions in *b*. *A*, which is called signature matrix, is the gene expression profile of cells.

In the first version of CIBERSORT, a machine learning technique, Nu-Support Vector Regression (*ν*-SVR), is used to solve the problem 1^47^. Matrix *A* in equation (1) is determined by a hyperplane with capturing the data points inside an *ε*-tube that is determined by support vectors (genes in signature matrix). SVR penalizes the data points outside the *ε*-tube, and a small value is used for *ν* that determines the lower bound of support vectors and the upper bound of training errors. Regression coefficients of *ν*-SVR method are the values of the vector *X*. However, the proportions are non-negative values, and their sum must be one. Therefore, negative coefficients are set to 0, and they normalize the remaining coefficients to sum to 1^47^. Newman et al.^18^ have recently improved their method by adding batch correction modes to remove possible cross-platform variations between signature matrix and mixture data.

To investigate the immune variations in renal cancer, we downloaded TCGA data set of gene expression profiles of 607 ccRCC primary tumors from UCSC Xena to use as a mixture data *b*. We used LM22 signature matrix, which includes 547 genes differentiating 22 cell types that are naive B cells, memory B cells, Plasma cells, CD8+ T-cells, CD4+ naive T-cells, CD4+ memory resting T-cells, CD4+ memory activated T-cells, follicular helper T-cells, regulatory T-cells (Tregs), *γδ* T-cells, resting NK cells, activated NK cells, monocytes, M0 macrophages, M1 macrophages, M2 macrophages, resting dendritic cells, activated dendritic cells, resting mast cells, activated mast cells, eosinophils, neutrophils. We then estimated cell fractions in ccRCC tumors using CIBERSORTx B-mode to remove technical differences between LM22 signature matrix and TCGA RNA-seq data.

After we estimated cell proportions, we included only cases with CIBERSORTx P-value < 0.05. We then applied unsupervised K-mean clustering algorithm to cluster tumors based on their percentage of immune cells. The K-mean algorithm separates samples in k-group of equal variance by minimizing the inertia (distance between samples in the clusters and center of the clusters). To determine the optimal number of clusters (k-value), we used elbow method to find the best value for *k*^48^.

We also collected clinical information of patients from cBioPortal and dropped some patients due to missing clinical information and continued our analysis with 526 patients. Patients’ characteristic are given in Table 1.

**Table 1.**
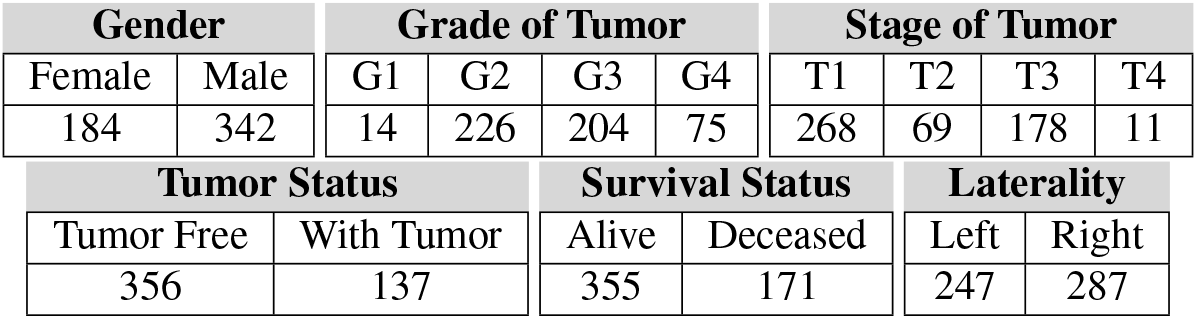
Patients’ characteristics. Sub-tables indicate the number of patients in each category. Differences in the numbers are due to missing information for some patients.

| Gender |  | Grade of Tumor |  |  |  | Stage of Tumor |  |  |  |
| --- | --- | --- | --- | --- | --- | --- | --- | --- | --- |
| Female | Male | G1 | G2 | G3 | G4 | T1 | T2 | T3 | T4 |
| 184 | 342 | 14 | 226 | 204 | 75 | 268 | 69 | 178 | 11 |
| Tumor Status |  | Survival Status |  | Laterality |  |  |  |  |  |
| Tumor Free | With Tumor | Alive | Deceased | Left | Right |  |  |  |  |
| 356 | 137 | 355 | 171 | 247 | 287 |  |  |  |  |

For statistical analyses, we used the non-parametric Mann-Whitney-Wilcoxon (MWW) test, because values in the comparison groups are not normally distributed and there are different numbers of patients in the comparison groups. MWW tests whether the values in one of two comparison groups is significantly larger than the other^49^. Stars in the figures show the significance levels where, ns: 0.05 < *p* ≤ 1, *: 0.01 < *p* ≤ 0.05, **: 0.001 < *p* ≤ 0.01, ***: 0.0001 < *p* ≤ 0.001, ****:*p ≤* 0.0001.

## Acknowledgements

Research reported in this publication was supported by the National Cancer Institute of the National Institutes of Health under Award Number R21CA242933.

## Author contributions statement

S.S. performed the analysis, analysed the results, and wrote the manuscript. S.A. prepared the data. L.S. designed and supervised the project, and participated in writing the manuscript. All authors reviewed the manuscript.

## Data availability

The data underlying this article are available at https://www.cbioportal.org/datasets and https://xenabrowser.net/datapages/.

## Ethics declarations

### Ethics

No ethical approval was required for this study.

